# Circuit dissection and functional validation of a cross-species emotional biomarker

**DOI:** 10.1101/2022.11.11.516218

**Authors:** Adam Jackson, Joshua Cohen, Aarron Phensy, Edward Chang, Heather Dawes, Vikaas S. Sohal

**Affiliations:** Department of Psychiatry and Behavioral Sciences, San Francisco, CA 94143-0444; Department of Neurological Surgery, San Francisco, CA 94143-0444; Weill Institute for Neurosciences, Center for Integrative Neuroscience, and Kavli Institute for Fundamental Neuroscience, University of California, San Francisco, San Francisco, CA 94143-0444

## Abstract

Emotional responses arise from limbic circuits including the hippocampus and amygdala. In the human brain, beta-frequency communication between these structures correlates with self-reported mood and anxiety. However, both the mechanism and significance of this biomarker as a readout vs. driver of emotional state remain unknown. Here we show that beta-frequency communication between the ventral hippocampus and basolateral amygdala also predicts anxiety-related behavior in mice on both long timescales (∼30 min) and immediately preceding behavioral choices. Genetically encoded voltage indicators reveal that this biomarker reflects synchronization between somatostatin interneurons across both structures. Indeed, synchrony between these neurons dynamically predicts approach vs. avoidance, and optogenetically shifting this synchronization by just 25 msec is sufficient to bidirectionally modulate anxiety-related behaviors. Thus, back-translation establishes a human biomarker as a causal determinant (not just predictor) of emotional state, revealing a novel mechanism whereby interregional synchronization that is frequency-, phase- and cell type-specific controls anxiety processing.

## INTRODUCTION

Under normal conditions, anxiety acts as an instructive signal to promote adaptive avoidance behaviors which assist individuals navigating through an uncertain world. However, these signals can also become maladaptive, producing excessive and inappropriate levels of avoidance characteristic of many anxiety disorders. Collectively, anxiety disorders comprise the most common psychiatric disorder in the world and result in a high burden of disease for those affected (Santomauro et al., 2021; Yang et al., 2021). The identification of robust and valid biomarkers of psychiatric disorders is a crucial tool for improving their prevention, diagnosis and treatment. Both preclinical and clinical studies have identified limbic structures as key regulators of emotion (LeDoux, 2000). Recent work using intracranial electroencephalogram (iEEG) recordings in humans identified an electrophysiological biomarker that tracks real-time fluctuations in an individual’s self-reported emotional state (Kirkby et al., 2018). This biomarker, which was based on beta-frequency coherence (β; 13-30Hz) between the amygdala (AMY) and hippocampus (HPC), represented a major advance in our understanding of how rapid timescale patterns of activity within the human brain encode emotional states. In particular, this biomarker was conserved across most subjects, could predict ∼40% of the variance in scores on a “Immediate Mood Scaler”, which includes items related to both mood and anxiety, and tended to be valid in individuals with high baseline levels of anxiety. However, this biomarker was identified within human iEEG recordings, making it challenging to investigate its underlying cellular mechanisms and functional significance more deeply.

Many preclinical studies have investigated the roles of the amygdala and hippocampus in anxiety-related behaviors in rodents. Lesions to the ventral hippocampus (vHC) reduce anxiety-related behaviors (Bannerman et al., 2003; Kjelstrup et al., 2002). Consistent with this, vHC is enriched with cells that are activated in anxiogenic environments, and opto-or chemogenetically modulating vHC outputs can bidirectionally influence anxiety-dependent behaviors (Ciocchi et al., 2015; Jimenez et al., 2018; Kheirbek et al., 2013; Padilla-Coreano et al., 2016; Parfitt et al., 2017; Sánchez-Bellot et al., 2022). The amygdala is composed of several subnuclei, of which the basolateral amygdala complex (BLA) and central amygdala have been implicated in anxiety processing (Babaev et al., 2018; Janak & Tye, 2015).

The BLA has strong projections which are thought to convey anxiety related information to the vHC (AlSubaie et al., 2021; Felix-Ortiz et al., 2013; McDonald & Mott, 2017; Pi et al., 2020). While these studies have highlighted a role of BLA-vHC projections in the regulation of anxiety behaviors, little is known about specific patterns of neural activity through which these areas functionally interact during risk assessment behaviors. Furthermore, it remains unknown whether such interactions give rise to biomarkers that can predict an individuals’ emotional state and corresponding behaviors in real-time.

Preclinical studies typically start with behaviors that are presumed (based on face validity) to reflect disease-relevant emotional states, then identify underlying patterns of neural activity, and implicitly assume that similar neural patterns will translate to the human condition. By contrast, here we started with an established human biomarker (amygdala-hippocampal β-coherence). Using homecage local field potential (LFP) recordings, we found that this human-derived biomarker also predicts the emotional states of mice (as reflected by their anxiety-related avoidance behavior). This biomarker was specifically associated with synchronization of somatostatin interneurons (SST+) within the BLA and vHC. Synchronization of these cell populations increases in anxiogenic environments and immediately prior to avoidance behaviors in the elevated plus maze. Artificially inducing or disrupting this synchronization, using 20 Hz stimulation that is either in-phase or shifted 25 msec out-of-phase, bidirectionally modulates these anxiety-related behaviors. Thus, this cross-species biomarker is a driver (not just a readout) of emotional state. Furthermore, these results reveal a previously unknown role for precise phase relationships between specific neuronal populations distributed across multiple limbic structures in creating network states which control emotional responses.

## RESULTS

### Hippocampus-amygdala β-coherence is a cross-species biomarker of negative emotion

We sought to test whether the variance of β-coherence between the amygdala and hippocampus, which tracks human mood and anxiety (Kirkby et al., 2018), also predicts anxiety levels in rodents. For this, we recorded LFP signals and computed the variance of β-coherence between the BLA and vHC in an animal’s homecage. Immediately after this recording session, we assayed behavior over 5min in the elevated zero maze (EZM). This testing was repeated on 3 days, separated by 48hr intervals (Figure 1A). As hypothesized based on human data, time spent exploring the open arms of the EZM showed a strong negative relationship with the variance of vHC-BLA β-coherence measured in the homecage (Figure 1B). Thus, increases in this biomarker indicate higher baseline anxiety levels. This correlation with behavior was not observed for the mean level of β-coherence or the mean or variance of coherence in other frequency bands (theta: 4-12 Hz, low gamma: 30-50 Hz, high gamma: 65-90 Hz) (Figure S1).

**Figure 1.**
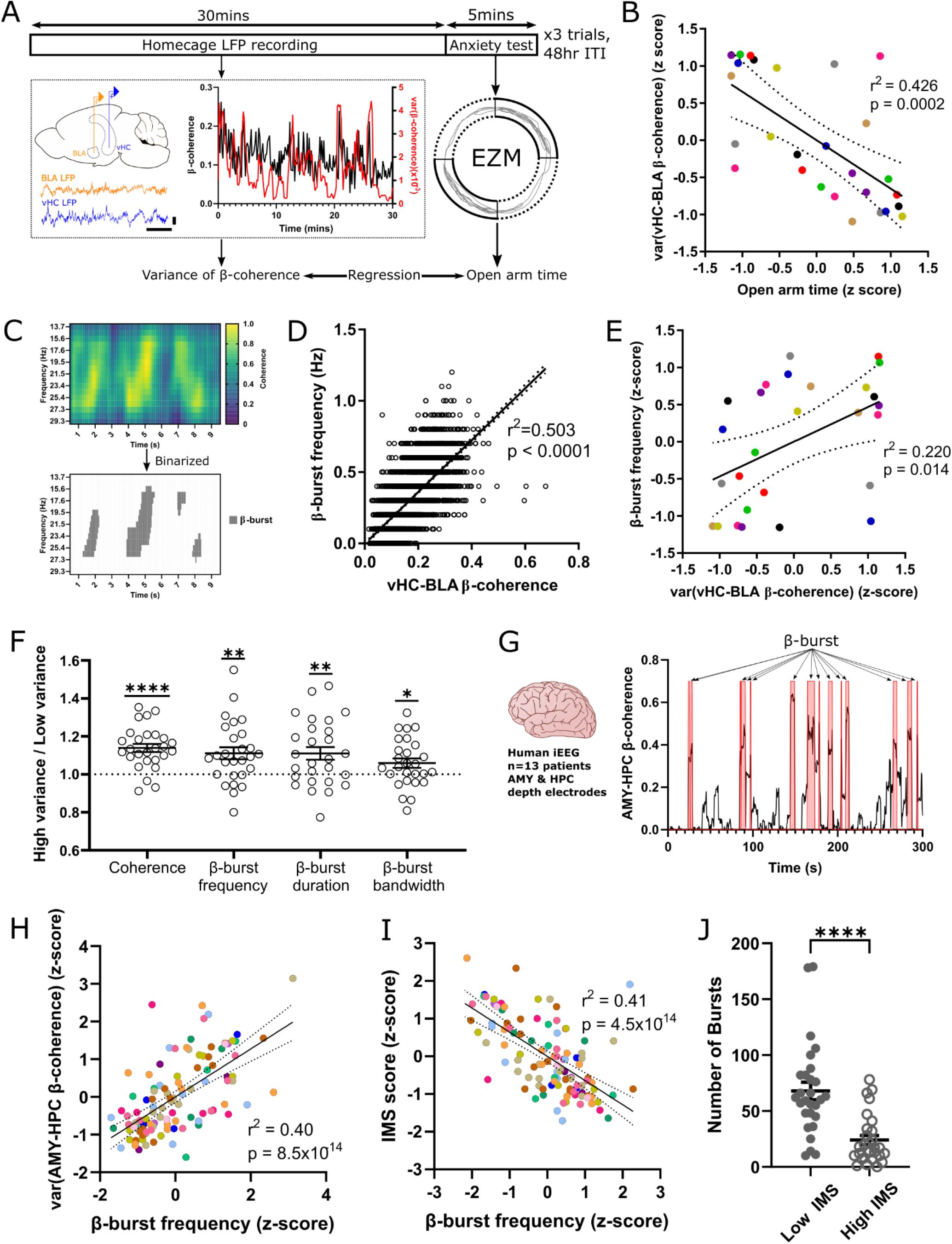
vHC-BLA variance of β-coherence is predictive of anxiety levels and is driven by transient high coherence bursts in both mice and humans. (A) Schematic of experimental design. Top: 30min homecage LFP recordings were analyzed and compared with immediate subsequent elevated zero maze behavior. Each mouse was recorded over 3 sessions with a 48hr inter-session interval. Bottom, left: LFP signals were recorded from electrodes implanted in BLA and vHC when animals were in their homecage. Bottom, center: vHC-BLA β-coherence was calculated across 10s non-overlapping windows (black) and coherence variance calculated over a 60s sliding interval (red). Bottom, right: Location tracking in EZM (grey). Time in open arms (broken lines) was calculated. (B) Regression of EZM open arm time against the variance of vHC-BLA β-coherence in homecage LFP recordings. Measures were z-scored within each subject. Dotted boundaries represent 95% confidence on the regression estimate. (C) Top: Example coherogram displaying brief periods of high β-coherence between vHC and BLA LFP signals. Bottom: Thresholded / binarized coherogram highlighting the presence of vHC-BLA β-bursts. (D) Regression of β-burst frequency against mean vHC-BLA β-coherence for 10s LFP recording windows. Dotted boundaries represent 95% confidence on the regression estimate. (E) Regression of variance of vHC-BLA β-coherence against β-burst frequency for each recording session. Measures were z-scored within each subject. Dotted boundaries represent 95% confidence on the regression estimate. (F) Mean coherence, β-burst frequency, duration and bandwidth are significantly greater during periods when the vHC-BLA β-coherence is high, compared to low variance periods (Coherence: t(26)=6.484, p<0.0001, β-burst frequency: t(26)=3.53, p=0.0016, β-burst duration: t(26)=3.3, p=0.0028, β-burst bandwidth: t(26)=2.426, p=0.0025; one-sample t test, n=27 sessions). (G) Example trace of AMY-HPC β-coherence (black) in human iEEG recordings, with β-bursts highlighted (red). (H) Regression of variance of AMY-HPC β-coherence against the frequency of β-bursts in human iEEG recordings. Both measures were z-scored within each subject. Dotted boundaries represent 95% confidence on the regression estimate. (I) Regression of variance of AMY-HPC β-coherence against the frequency of β-bursts. All measures were calculated over a 20-min window centered around each IMS score time point and z-scored within each subject. Dotted boundaries represent 95% confidence on the regression estimate. (J) β-burst frequency was significantly increased during periods of low IMS score compared to high IMS periods (p<0.0001, Low mood n=28, High mood n=28; Mann-Whitney test).

The variance of coherence was calculated using a 60s window, suggesting that high variance may reflect transient periods of high coherence occurring on faster timescales and superimposed on a baseline of lower coherence. Indeed, time-frequency plots for vHC-BLA β-coherence reveal brief periods of high β-coherence, termed here ‘β-bursts’ (Figure 1C). The frequency of these bursts strongly predicts the overall level of vHC-BLA β-coherence within each 10s window (Figure 1D), and also had a significant positive relationship with β-coherence variance (Figure 1E), suggesting that β-bursts are a major contributor to this biomarker. Comparing periods of β-coherence with high variance (upper quartile) to low variance periods (bottom quartile) reveals that high β-coherence variance is associated with increases in the number, duration and bandwidth of β-bursts (Figure 1F). Similar periods of brief, high coherence were also observed for AMY-HPC β-coherence in human intracranial electroencephalography (iEEG) recordings (Figure 1H). In human data, the frequency of β-bursts had a strong, positive correlation with the variance in AMY-HPC β-coherence biomarker (Figure 1I). Additionally, β-burst frequency was strongly negatively correlated with subjective mood (Figure 1J). Correspondingly, recordings during periods of low mood (bottom quartile) contained significantly more β-bursts than high mood periods (upper quartile) (Figure 1K).

### β-bursts predict anxiety-related behaviors on fast timescales

Thus the variance of hippocampus-amygdala β-coherence, which is biomarker for emotional state in both humans and mice, is, in both species, driven (at least in part) by β-bursts occurring on much faster timescales. Whereas coherence variance is calculated over 1 min windows and then averaged over tens of minutes, β-bursts occur on timescales of ∼1s, enabling us to ask whether they play a role in the realtime production of anxiety-related behaviors on these rapid, behaviorally-relevant timescales. To assess this, we used the elevated plus maze (EPM), which contains a center zone in which mice choose to either approach an exposed and brightly-lit open arm (open runs; green arrow) or engage in anxiety-related avoidance by entering a closed arm (closed runs; black arrow) (Figure 2A). We observed a time-locked increase in β-burst probability immediately prior to avoidance behavior that was notably absent when animals approached the open arm (Figure 2B). This was reflected by a higher β-burst frequency during the 3s prior to closed arm entry compared to open arm entry (Figure 2C). Although the average speed of locomotion was higher during closed runs than open runs, the average locomotion speed when mice were still in the center zone (before they entered the closed or open arms) did not differ for closed vs. open runs (Figure S2). Furthermore, there was no relationship between locomotion speed and β-burst frequency for either closed arm or open arm runs (Figure S2). Thus, the increase in β-burst frequency associated with open arm avoidance was not simply an artifact of differences in locomotion speed.

**Figure 2.**
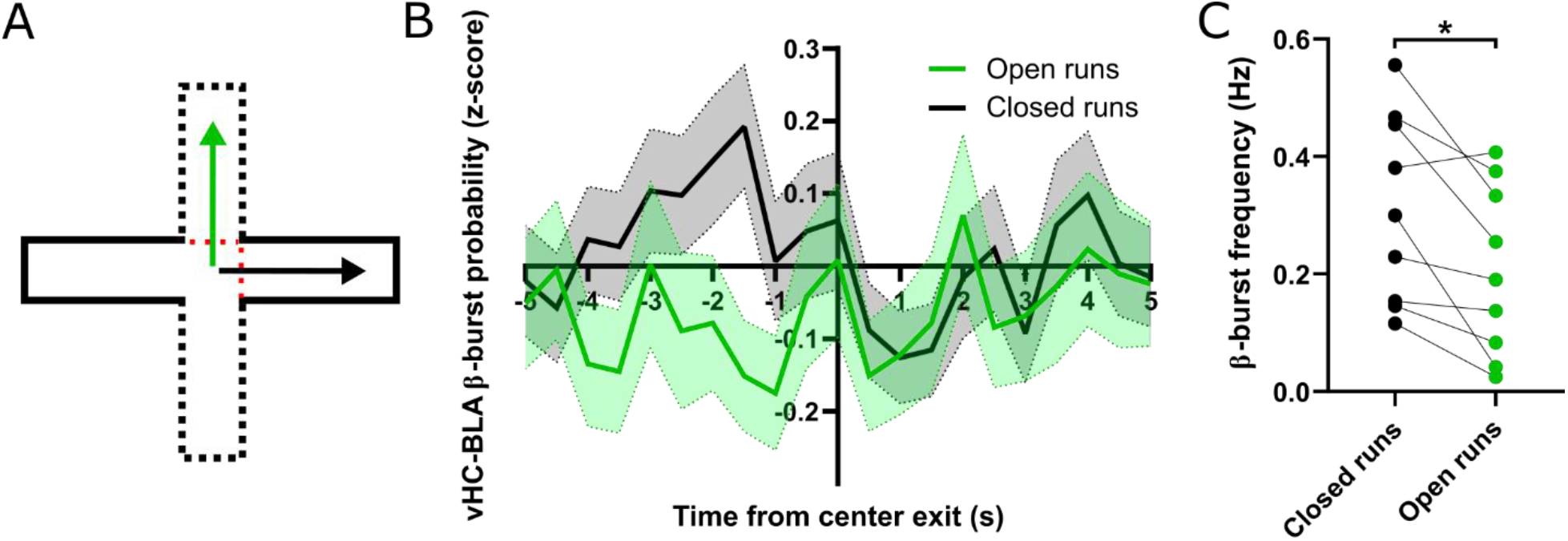
vHC-BLA β-bursts predict future exploration versus avoidance. (A) Experimental design schematic. LFP signals were analyzed, time-locked to when mice exited the center of the EPM during entries to either the open (green arrow) or closed arm (black arrow). (B) Time course of vHC-BLA β-burst probability as animal engage in anxiety-based approach (open arm entries) or avoidance (closed arm entries) behavior. (C) vHC-BLA β-burst frequency was significantly higher during the 3s prior to avoidance behavior than when animals approached the open arm (p=0.0122, n=9 mice; paired t test).

### vHC and BLA somatostatin interneurons synchronize during β-bursts

GABAergic interneurons have been implicated in generating and maintaining neuronal oscillations throughout the brain. We next tested whether β-bursts were accompanied by the synchronization of specific GABAergic cell types within the BLA and vHC using trans-membrane electrical measurements performed optically (TEMPO). TEMPO uses bulk fluorescence from the voltage indicator Ace2N-4AA-mNeon (mNeon) to monitor voltage signals in specific cell types (Marshall et al., 2016). We injected AAV-CAG-DIO-Ace2N-4AA-mNeon and AAV-Synapsin-tdTomato unilaterally into both the BLA and the vHC of *SST-Cre* or *PV-Cre* mice to drive Cre-dependent expression of mNeon in SST+ or PV+ interneurons respectively, along with nonspecific expression of a reference fluorophore (tdTomato). We then implanted optrodes to measure mNeon and tdTomato fluorescence as well as LFP in these areas (Figure 3A) while animals were in their homecage (Figure 3B). Fluorescence from genetically encoded voltage indicators is contaminated by both movement and hemodynamic artifacts (Carandini et al., 2015; Marshall et al., 2016). To minimize the influence of these artifacts in all analyses, after filtering mNeon and tdTomato signals in the frequency band of interest, we used robust linear regression to fit each tdTomato signal to the corresponding mNeon signal, then subtracted the rescaled tdTomato signal from the mNeon signal to obtain a partially corrected mNeon signal (Figure 3C). To assess the relationship between interneuron synchrony and vHC-BLA β-bursts, the onset of each β-burst was identified in LFP recordings and time-locked to the simultaneously recorded mNeon signals (filtered at 13-30Hz) (Figure 3D). Correlations between vHC and BLA mNeon signals were assessed using a 250ms sliding window with an 80% overlap. For each timepoint, we also compared the correlation in real data to 100 shuffled data sets that correlated one real mNeon signal with a randomly time-shifted mNeon signal. In *SST-Cre* mice, 13-30Hz filtered mNeon signals showed an increase in correlation around the onset of β-bursts.

**Figure 3.**
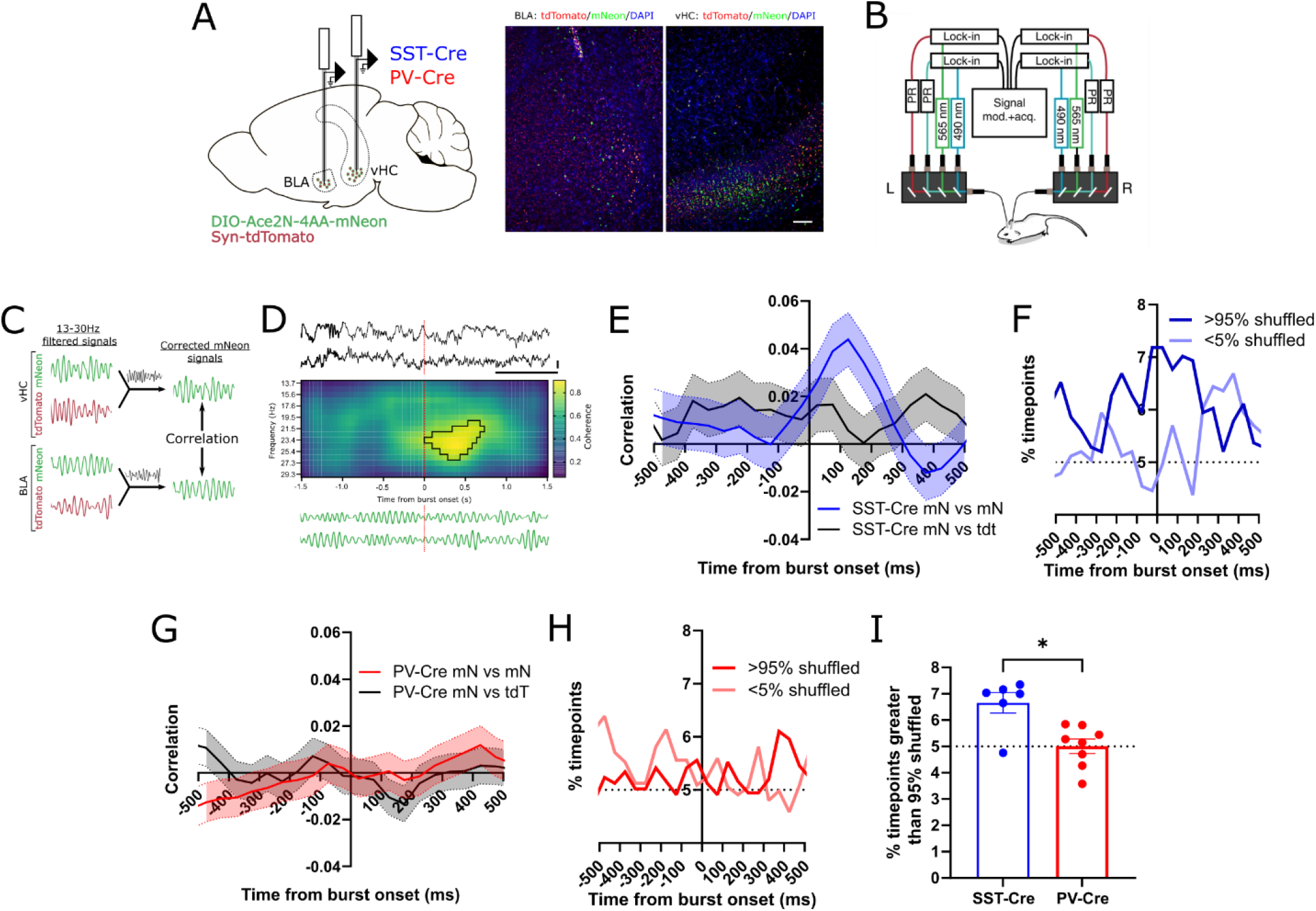
vHC and BLA Somatostatin positive interneurons synchronize during vHC-BLA β-bursts. (A) Schematic illustrating the experimental design. *SST-Cre* or *PV-Cre* mice had unilateral injections of AAV-DIO-Ace2N-4AA-mNeon and AAV-Syn-tdTomato injections plus optrode implants in the BLA and vHC. Scale bar: 100μm. (B) Schematic of dual-site TEMPO set up. Each fiber-optic implant, for delivering illumination and collecting fluorescence, connects to a mini-cube coupled to two LEDs and two photoreceivers (PRs) to separately excite and collect emitted fluorescence from Ace–mNeon and tdTomato. Two lock-in amplifiers modulate (mod.) LED output and demodulate PR signals, which are then acquired (acq.) by a multichannel real-time signal processor. (C) Overview of TEMPO analysis: mNeon and tdTomato signals from each region were filtered around the frequency of interest. To minimize the influence of hemodynamic and movement artifacts within mNeon signals, the mNeon signal from each region was also ‘corrected’ by subtracting the robust linear regression fit of the tdTomato signal to that mNeon signal. (D) Correlation between mNeon signals from both regions were analyzed around the onset of vHC-BLA β-bursts within LFP recordings. Correlations were compared to time-shuffled mNeon data. Scale bar: 200μV, 500ms. (E) SST+ interneuron 13-30Hz mNeon correlation values aligned to the onset of vHC-BLA β-bursts. For comparison, we show correlation between mNeon signals in the two regions (blue) and the correlation between mNeon signals and the tdTomato signal in the other region (black). (F) Trace showing the fraction of SST+ interneuron mNeon correlation values which exceed / fall below the 95^th^ / 5^th^ percentiles based on time-shuffled mNeon signals. (G) PV+ interneuron 13-30Hz mNeon correlation values aligned to the onset of vHC-BLA β-bursts. For comparison, we show correlation between mNeon signals in the two regions (red) and the correlation between mNeon signals and the tdTomato signal in the other region (black). (H) Trace showing the fraction of PV+ interneuron mNeon correlation values which exceed / fall below the 95^th^ / 5^th^ percentiles based on time-shuffled mNeon signals. (I) β-synchrony in SST+ interneurons was significantly enhanced compared to PV+ interneurons during the first 250ms of vHC-BLA β-bursts (p=0.0127, *SST-Cre* n=6, *PV-Cre* n=8; Mann-Whitney test).

Notably, there was no increase in correlation between each region’s mNeon signal and the non-voltage dependent tdTomato signal from the other region, confirming that this increased correlation was not driven by artifacts (Figure 3E&F). We also did not observe increased 13-30Hz synchrony (i.e., an increase in the correlation of filtered mNeon signals) around the onset of β-bursts in *PV-Cre* mice (Figure 3G&H). Correspondingly, during the first 250ms of β-bursts, the percentage of timepoints with correlations greater in real data than in 95% of the shuffled datasets was significantly greater for *SST-Cre* than *PV-Cre* mice (Figure 3I). By contrast, no changes in gamma-frequency synchrony (i.e., the correlation between 30-50 Hz filtered mNeon signals) were observed around the onset of β-bursts for either SST+ or PV+ interneurons (Figure S3).

### vHC and BLA SST+ interneuron synchrony is elevated during anxiety-dependent behavior

Given this relationship between the β-synchrony of vHC and BLA SST+ interneurons and the β-bursts which precede anxiety-related avoidance behaviors, we next tested whether SST+ interneuron β-synchrony was similarly modulated by an animal’s anxiety state. SST+ interneuron β-synchrony increased during EPM sessions relative to same day homecage recordings (Figure 4A). This effect was driven by a specific increase in synchrony when the animal was in the center zone of the EPM (Figure 4B). Neither this general increase in β-synchrony in the EPM (Figure 4C) nor any more specific changes within individual zones of the EPM (Figure 4D) were observed in PV+ interneurons. These effects were specific for the β-frequency range and not observed for either SST+ or PV+ interneurons when mNeon signals were filtered from 30-50Hz (Figure S4).

**Figure 4.**
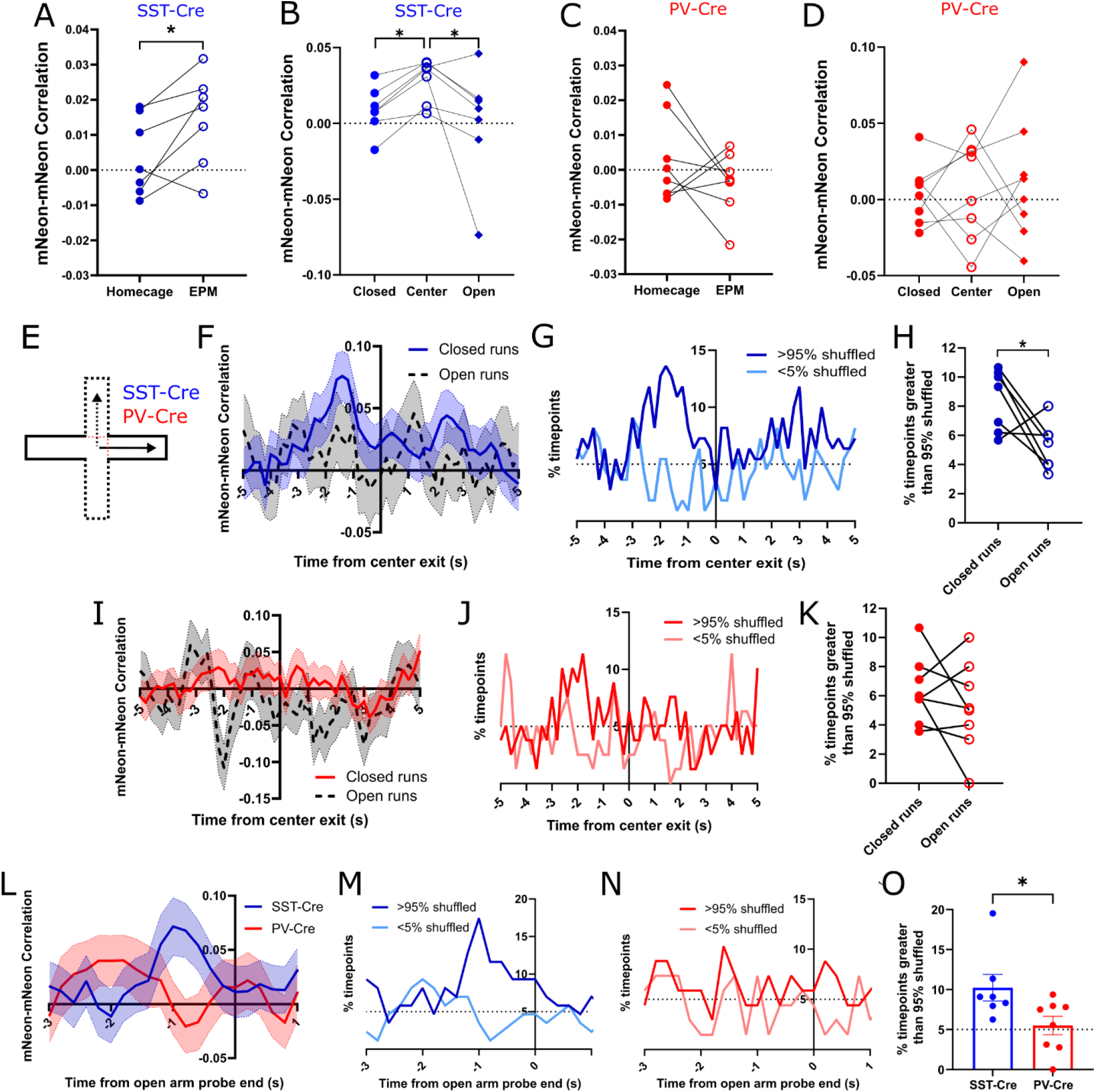
vHC and BLA SST+ interneurons reflect and predict elevated plus maze behavior. (A) SST+ interneuron β-synchrony was significantly higher during sessions of EPM exploration compared to the homecage (p=0.038, n=7 mice; paired t test). (B) SST+ interneuron β-synchrony was significantly increased in the EPM center zone relative to the open and closed arms (p = 0.0207, n=7 mice; Friedman test with Dunn’s correction; closed vs center: post hoc p=0.0485, closed vs open: post hoc p>0.99, center vs open: post hoc p=0.0485). (C) PV+ interneuron β-synchrony in the homecage vs. EPM (p=0.2985, n=8 mice; paired t test). (D) PV+ interneuron β-synchrony was not consistently influenced by an animal’s location within the EPM (p = 0.7829, n=8 mice; Repeated measures one-way ANOVA). (E) Schematic of experimental design. The timecourse of TEMPO signals were analyzed, time-locked to exits from the center zone, depending on whether mice subsequently entered either the open (dashed arrow) or closed arms (solid arrow). (F) SST+ interneuron 13-30Hz mNeon correlation values aligned to closed vs. open arm entry. (G) Trace showing the fraction of SST+ interneuron mNeon correlation values which exceed / fall below the 95^th^ / 5^th^ percentiles based on time-shuffled mNeon signals. (H) SST+ interneurons β-synchrony was significantly greater during 5s prior to closed arm entries compared to open arm entries (p=0.039, n=7 mice; paired t test). (I) PV+ interneuron 13-30Hz mNeon correlation values aligned to closed and open arm entry. (J) Trace showing the fraction of PV+ interneuron mNeon correlation values which exceed / fall below the 95^th^ / 5^th^ percentiles based on time-shuffled mNeon signals. (K) PV+ interneuron β-synchrony during the 5s prior to closed or open arm entries (p=0.3557, n=8 mice; paired t test). (L) SST+ and PV+ interneuron 13-30Hz mNeon correlation values aligned to the end of open arm probes. (M) Time course of SST+ interneuron mNeon correlation values during open arm probes, expressed as the fraction of values which exceed the 95th / are below the 5th percentile of time-shuffled mNeon signals. (N) Time course of PV+ interneuron mNeon correlation values during open arm probes, expressed as the fraction of values which exceed the 95th / are below the 5th percentile of time-shuffled mNeon signals. (O) β-synchrony was significantly greater for SST+ interneurons than PV+ interneurons during the 1.5s prior to the termination of open arm probes (p=0.027, *SST-Cre* n=7, *PV-Cre* n=8; Mann-Whitney test).

Our LFP data indicated that β-bursts specifically precede open arm avoidance (Figure 2B). Therefore, we next examined whether the increase in SST+ interneuron β-synchrony we observed in the center zone of the EPM reflected a specific increase during avoidance behaviors. Signals time-locked to exit from the center zone revealed an increase in β-synchrony (the correlation between 13-30 Hz filtered mNeon signals) prior to open arm avoidance (i.e., closed arm entries), but not prior to open arm approaches (Figure 4F). The percentage of timepoints for which this correlation, calculated from real data, exceeded the 95^th^ percentile of values based on shuffled data also increased during the 5s immediately prior to open arm avoidance (Figure 4G) and was significantly greater than during the equivalent period preceding open arm approaches (Figure 4H). These effects were not observed in PV+ interneurons (Figure 4I-K) or in mNeon signals that had been filtered from 30-50Hz (Figure S4). For SST+ interneurons, but not PV+ interneurons, the correlation between 13-30Hz filtered mNeon signals also increased just prior to the termination of open arm “probes,” during which an animal’s head enters the open arm whilst its body remains in the center zone (Figure 4L). The fraction of timepoints for which the correlation between 13-30Hz filtered mNeon signals exceeded the 95^th^ percentile based on shuffled data also increased prior to the termination of open arm probes for SST+ interneurons (Figure 4M) but not PV+ interneurons (Figure 4N). Correspondingly, for the 1.5s preceding the termination of open arm probes, the fraction of timepoints for which the correlation between 13-30Hz filtered mNeon signals exceeded the 95^th^ percentile based on shuffled data was significantly greater in *SST-Cre* mice compared to *PV-Cre* mice (Figure 4O).

### Optogenetic induction or disruption of vHC-BLA SST+ interneuron β-synchrony bidirectionally modulates anxiety-related behaviors

Having established that β-synchrony of SST+ interneurons across the vHC and BLA is an anxiety-related biomarker, we next tested whether manipulating this pattern of circuit activity would causally influence anxiety-related risk-assessment behaviors. Bilateral fiber optics were implanted in the BLA and vHC of both *SST-Cre* and *PV-Cre* mice injected with Cre-dependent channelrhodopsin (AAV-DIO-ChR2-eYFP) (Figure 5A). As increases in β-synchrony occurred specifically within the center zone of the EPM, we delivered 473nm light whenever mice were in a ‘stimulation zone’ comprising the center zone and the proximal thirds of the open and closed arms. Behavior was compared across 3 different 3min epochs: a no stimulation period, an ‘in-phase’ period during which the same stimulation was delivered bilaterally to both the BLA and the vHC, and an ‘out-of-phase’ period during which bilateral stimulation was delivered that was 180° out of phase between the BLA and vHC (Figure 5B). Out-of-phase 20Hz stimulation in *SST-Cre* mice increased the amount of time mice spent in the center of the EPM relative to both the no stimulation and in-phase stimulation periods (Figure 5C). This effect (out-of-phase stimulation increases center time) was not observed when stimulation was delivered at 8Hz (instead of 20Hz) or in *PV-Cre* mice (instead of *SST-Cre* mice) (Figure 5C). Whilst no significant changes in open arm time were observed in any of the experimental groups (Figure 5D), out-of-phase 20Hz stimulation decreased the amount of time *SST-Cre* mice spend in the closed arm (Figure 5E). Within the center zone, stimulation in *SST-Cre* mice further modulated behavior in a phase and frequency-dependent manner.

**Figure 5.**
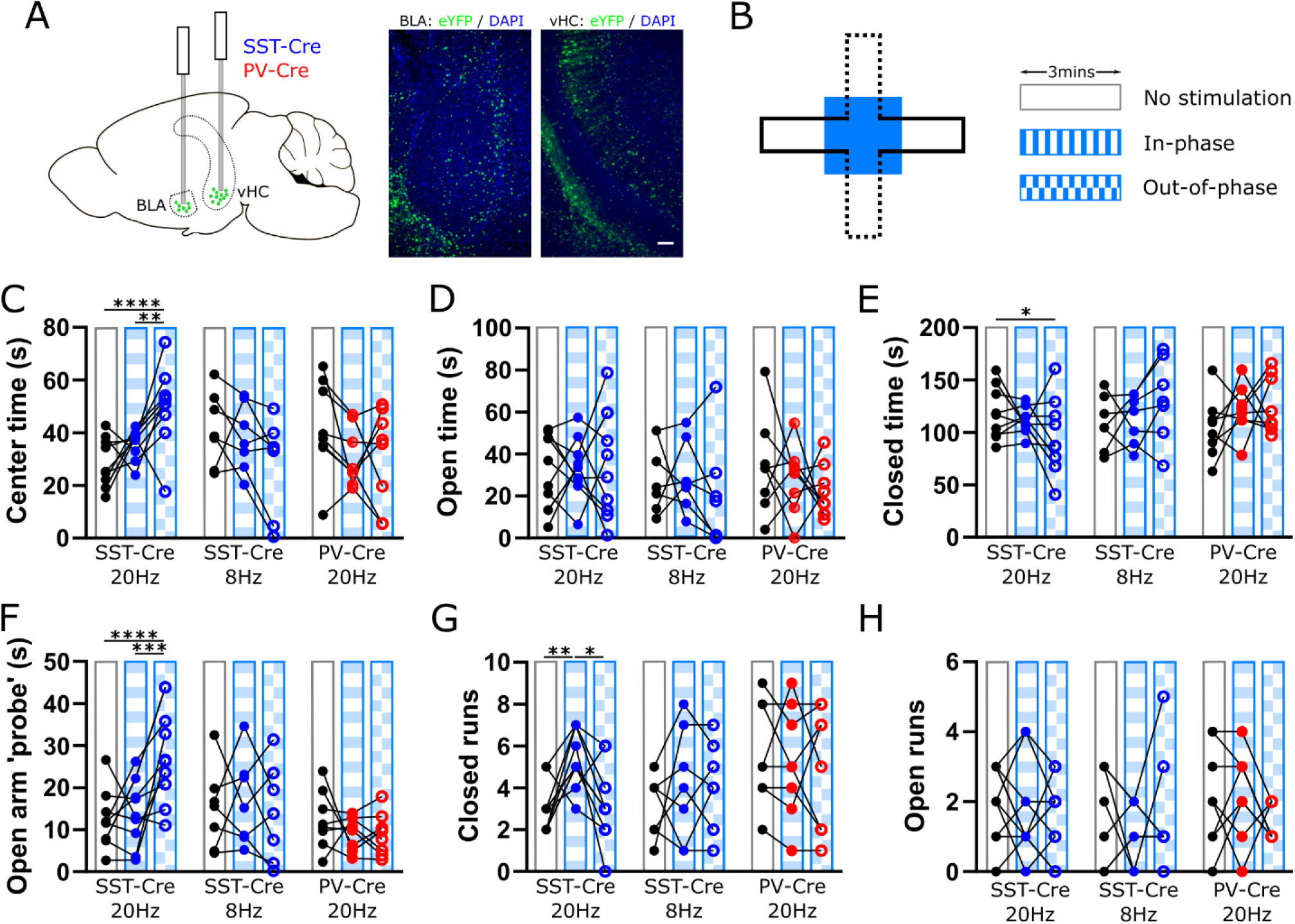
Modulating vHC-BLA SST+ interneuron β-synchrony bidirectionally influences approach-avoidance behaviors. (A) Experimental design: *SST-Cre* or *PV-Cre* mice received bilateral injections of AAV-DIO-ChR2 and optic fiber implants in the BLA and vHC. Scale bar: 100μm. (B) Stimulation paradigm: 473nm stimulation was restricted to the center zone and proximal portions of the open and closed arms in the EPM. Stimulation varied across three 3min epochs: no stimulation, in-phase stimulation and out-of-phase stimulation. (C) 20Hz out-of-phase stimulation in *SST-Cre* mice significantly increased time spent in the center zone (Two-way ANOVA (group x stim session); interaction: F_(4,40)_=7.848, p<0.0001; *SST-Cre* 20Hz post hoc with Tukey’s multiple comparison test: no stim vs in-phase: p=0.259; no stim vs out-of-phase: p=0.001; in-phase vs out-of-phase: p=0.037). (D) In-phase or out-of-phase stimulation at 8 or 20Hz had no effect on time in open arm in any group (Two-way ANOVA (group x stim session); interaction: F_(4,40)_=0.7748, p=0.548). (E) 20Hz out-of-phase stimulation in *SST-Cre* mice significantly decreased time spent in the closed arms (Two-way ANOVA (group x stim session); interaction: F_(4,40)_=3.435, p=0.017; *SST-Cre* 20Hz post hoc Tukey’s multiple comparison test: no stim vs in-phase: p=0.607; no stim vs out-of-phase: p=0.044; in-phase vs out-of-phase: p=0.288). (F) 20Hz out-of-phase stimulation in *SST-Cre* mice significantly increased open arm probing (Two-way ANOVA (group x stim session); interaction: F_(4,40)_=4.345, p=0.005; *SST-Cre* 20Hz post hoc Tukey’s multiple comparison test: no stim vs in-phase: p=0.712; no stim vs out-of-phase: p<0.0001; in-phase vs out-of-phase: p=0.001). (G) 20Hz in-phase stimulation in *SST-Cre* mice significantly increased the number of closed arm runs (Two-way ANOVA (group x stim session); interaction: F_(4,44)_=2.211, p=0.083; *SST-Cre* 20Hz post hoc Tukey’s multiple comparison test: no stim vs in-phase: p=0.009; no stim vs out-of-phase: p=0.946; in-phase vs out-of-phase: p=0.022). (H) In-phase or out-of-phase stimulation at 8Hz or 20Hz had no effect on open arm runs in any group (Two-way ANOVA (group x stim session); interaction: F_(4,44)_=0.868, p=0.491).

Specifically, out-of-phase 20Hz stimulation in *SST-Cre* mice increased the amount of open arm probing (Figure 5F). Again, this effect was not observed with 8Hz stimulation (in *SST-Cre*) mice or in *PV-Cre* mice (Figure 5F). In contrast to the effect of out-of-phase 20 Hz stimulation to increase center time and open arm probes in *SST-Cre* mice, in-phase 20Hz stimulation specifically increased closed arm runs in *SST-Cre* mice (Figure 5G). Neither in-phase or out-of-phase stimulation had any effect on the number of open arm runs in any of the experimental groups (Figure 5H). To confirm the role of anxiety in the observed effects, *SST-cre* mice performed the same testing within a cross maze, a modified EPM containing 4 closed arms. Neither in-phase nor out-of-phase 20Hz stimulation influenced center time in the cross maze (Figure S5). Additionally, neither stimulation pattern altered the time spent probing the cross maze arms that are geographically equivalent to either the open or closed arms in the EPM (Figure S5). This shows that the observed effects in the EPM are the result of anxiety-dependent approach-avoidance conflict, and not simply an artifact related to the geometry of the maze or non-specific effects of stimulation on behavior at choice points, etc.

## DISCUSSION

This study reveals that β-frequency communication between the amygdala and hippocampus predicts emotional state in mice, as was previously found in humans. Specifically, the variance of β-coherence between the two structures during homecage recordings correlates with subsequent avoidance behavior in the elevated zero maze. In addition to this cross-species translation, there are three other major advances. First, whereas the original description of this biomarker in humans and our initial back-translation to rodents measured this biomarker and its relationship to emotional state on timescales of minutes, we now find that in both humans and mice the biomarker is driven by ‘β-bursts’, transient periods of high β-coherence, which (in mice) predict approach-avoidance behaviors on timescales of seconds. Second, using genetically encoded voltage indicators, we found that these β-bursts were associated with β-frequency synchronization between SST+ interneurons (but not PV+ interneurons) in the BLA and vHC, and that the β-synchrony of these populations similarly increases during avoidance.

Finally, artificially inducing or disrupting synchrony of these SST+ populations (but not PV+ interneurons) during approach-avoidance conflicts can bidirectionally influence approach-avoidance behaviors in a phase and frequency dependent manner. Overall, this shows that amygdala-hippocampus β-frequency communication is a cross-species biomarker which depends on specific cell types and not only predicts emotional states, but also causally influences related behaviors.

### Precise phase-relationships define distinct anxiety-related network states

Prevailing paradigms in studies of circuits mediating emotional behaviors have largely focused on how changes in the activity of specific cell types (typically projection neurons with specific targets or GABAergic neurons expressing specific markers) on a timescale of seconds relative to specific behavioral events promotes or suppresses specific behaviors. An alternative has been to identify engrams, subsets of neurons that are activated during specific behaviors, e.g., freezing. This study reveals a novel and complementary dimension, whereby the synchronization of specific cell populations on much faster timescales (∼10-50 msec) and across brain regions defines network states which serve as markers of emotional processes that causally influence approach-avoidance behaviors. An important line of previous work has established an association between hippocampal-prefrontal theta-frequency synchronization and anxiety-related behavior (Adhikari et al., 2010, 2011; Kjaerby et al., 2016; Lee et al., 2019; Padilla-Coreano et al., 2016, 2019). These studies, particularly the finding that 8 Hz stimulation of hippocampal-prefrontal projections increases avoidance (Padilla-Coreano et al., 2019), were interpreted to indicate that anxiety-related input from the vHC to prefrontal cortex is normally modulated at theta-frequency, such that stimulation at this frequency is best for facilitating the transmission of this input. Here we find that two patterns of stimulation that are both delivered at the same frequency can nevertheless elicit opposing effects on approach-avoidance behaviors in a phase-dependent manner. This shows that it is the precise phase relationship between neurons in different brain structures, not just the strength of a rhythmic input or level of activity in a specific circuit, which defines distinct network states.

It is particularly notable that all of the effects we found, using both voltage indicators and optogenetics, were specific for cell type (present for SST+ interneurons absent for PV+ interneurons), frequency (changes in synchrony observed for mNeon signals filtered at 13-30Hz but not 30-50Hz, optogenetic manipulations affect behavior for 20Hz in/out of phase stim but not for 8Hz stim), and phase relationship (opposing behavioral effects of in-phase vs. 25 msec out-of-phase stim). These are powerful controls which confirm that these phenomena reflect the rhythmic synchronization of neuronal activity within specific cell types, not just overall increases in neuronal activity or non-neuronal artifacts. This new framework, wherein the phase relationships of interneuron populations across brain structures at specific frequencies defines network states which causally influences behavior, raises many important questions for future studies. First, how do these precise phase relationships arise? The BLA contains GABAergic neurons (including SST+ neurons) which project to both excitatory and inhibitory cells in the vHC (AlSubaie et al., 2021; Felix-Ortiz et al., 2013). The vHC also contains GABAergic neurons that project to several other subcortical and cortical targets (Jinno, 2009). This raises the possibility that direct connections exist between SST+ GABAergic neurons in the amygdala and hippocampus.

Second, how exactly does the presence of β-synchrony between interneuron populations in different (interconnected) brain structures affect information processing in ways that end up impacting approach-avoidance behaviors? The vHC contains multiple non-overlapping output streams that connect it with other nodes of the anxiety network, allowing it to bidirectionally influence anxiety states based on the differential recruitment of particular vHC projection neuron subnetworks (Ciocchi et al., 2015; Jimenez et al., 2018; Parfitt et al., 2017). The BLA is also hypothesized to transmit anxiety-related information to vHC circuits (AlSubaie et al., 2021; Felix-Ortiz et al., 2013; Pi et al., 2020). Thus, if β-frequency activity in SST+ interneurons entrains BLA output, then synchronization between SST+ inhibitory microcircuits in the BLA and vHC inputs may impact how the vHC integrates BLA inputs, thereby regulating the effective gain of these anxiety-related inputs and/or their propagation into specific classes of vHC projection neurons.

### Hippocampus-amygdala β-communication is a cross-species biomarker of anxiety

Previous studies that have attempted to identify neural signatures of anxiety in rodents have done so by comparing brain activity when mice are in areas that are assumed to be safe versus anxiogenic, or examining genetic models that exhibit pathological anxiety (Adhikari et al., 2011; Cruces-Solis et al., 2021; Cunniff et al., 2020; Jacinto et al., 2016). By contrast, here we used neural recordings from wild type mice in a familiar, homecage environment to predict their current anxiety level, which was then assayed by subsequent exposure to the EZM. In both human and mouse recordings, we find that increases in β-coherence variance are driven by transient bursts of high β-coherence. β-frequency activity in somatosensory and motor regions has previously been shown to consist of similar transient high-power β-events (Feingold et al., 2015; Sherman et al., 2016; Shin et al., 2017). β-frequency events have also been shown to influence sensory perception in both mice and humans (Shin et al., 2017), further highlighting the broad relevance of this pattern of circuit dynamics across species and brain regions. Furthermore, whilst there is not a 1:1 relationship between interneuron subtypes and specific oscillation frequencies, SST+ interneurons have previously been implicated in oscillations at the upper end of the β-frequency / lower end of the gamma-frequency range (Chen et al., 2017; Veit et al., 2017).

Whilst β-frequency oscillations are traditionally associated with sensorimotor processing, increasing evidence has implicated them in a variety of cognitive functions including decision making, working memory and reward processing (Lundqvist et al., 2016; Spitzer & Haegens, 2017). Recently, β-coherence within the anxiety network, including vHC-BLA interactions, was found to be increased in a mouse model with increased anxiety levels (Cruces-Solis et al., 2021) suggesting that β-synchrony may contribute to pathological as well as typical or adaptive manifestations of anxiety. An interesting future direction will be to investigate how anxiolytic drugs or perturbations that exacerbate anxiety affect this pattern of limbic circuit activity.

A simple view of the EPM is that it comprises ‘safe’ closed arms, ‘anxiogenic’ open arms, and a center zone that is intermediate, both in terms of its location and its relationship to anxiety. One problem with this is the ambiguity about whether open arm times should be regarded as periods when the mouse is less anxious (because it is willing to explore the open arms) or more anxious (because it is exposed to the open arms)? In this context, it is notable that we found that the β-synchrony of SST+ neurons across the BLA and vHC was actually higher in the center zone than both the closed and open arms, and specifically increases just before decisions to avoid the open arms. This suggests that vHC-BLA SST+ neuron β-synchrony reflects the specific approach-avoidance conflict which occurs in the center zone, rather than more nebulous and anthropomorphic notions of ‘anxiety’ traditionally associated with the open arms.

### Optogenetic modulation of synchrony bidirectionally influences anxiety-related behaviors

While identifying a correlative biomarker can be useful for diagnosis, treatment stratification, or emotional readout, it does not address whether that biomarker is a causal driver of associated behaviors. Here, using optogenetics, we were able to show that β-synchrony between the BLA and vHC bidirectionally influences anxiety-related behaviors in a cell type, phase and frequency specific manner. Artificially inducing β-synchrony via in-phase stimulation was sufficient to promote anxiety-related avoidance (closed arm runs). Conversely, out-of-phase stimulation that desynchronizes the same populations was able to increase risk assessment behaviors (center zone time and open arm probes), although it was not on its own sufficient to induce the exploration of normally anxiogenic regions. These results indicate that approach behaviors in the EPM can be decomposed into at least two stages representing ‘risk assessment’ (remaining in the center zone, performing open arm probes) and ‘exploration’ (runs into the open arms), respectively. In this formulation, desynchronizing out-of-phase stimulation is sufficient to promote the first, but not the second, of these stages. This furthermore aligns with our observation that vHC-BLA SST+ neuron β-synchrony is higher in the center zone than in both the open and closed arms. Thus, this pattern of synchronization seems to reflect and causally influence risk-assessment (in the center zone) rather than the drive to explore the open arms.

## Conclusion

β-synchrony between the amygdala and hippocampus is a cross-species biomarker of emotional state on both long and short timescales. This biomarker is driven by the synchrony of SST+ interneuron populations within these structures. Altering the inter-regional phase relationships of these neurons is sufficient to bidirectionally influence avoidance and risk assessment behaviors. These findings reveal underlying cellular substrates and behavioral functions for patterns of activity that have previously been linked to emotional state in the human brain. They further establish phase relationships between specific neuronal populations in different brain structures as a key determinant of network state encoding emotion. This both informs our basic understanding of how the brain encodes emotion, and may lead to new interventions which modify network states defined by synchrony between specific cell types in order to treat pathological forms of anxiety.

## ACKNOWLEDGEMENTS

This work was supported by the UCSF Dolby Family Center for Mood Disorders and NIMH (R01 MH117961 to VSS).

## METHODS

### Animals

All animal care procedures and experiments were conducted in accordance with the National Institutes of Health guidelines and approved by the Administrative Panels on Laboratory Animal Care at the University of California, San Francisco. Mice were housed in a temperature-controlled environment (22– 24 °C) with ad libitum access to food and water. Mice were reared in normal lighting conditions (12-h light/dark cycle). Mice aged 10-16 weeks of the following lines were used: Wild-type C57Bl/6 mice, Sst < tm2.1(cre)Zjh > /J (line 013044; https://www.jax.org), and B6;129P2-Pvalb < tm1(cre)Arbr > /J (line 008069; https://www.jax.org).

### General Surgery

Mice were anaesthetized using isoflurane (2.5% induction, 1-1.5% maintenance in 95% oxygen, flow rate 1L/min) and secured using ear bars in a stereotaxic frame (David Kopf instruments). Body temperature was maintained using a heating pad. The scalp and the periosteum were removed from the dorsal surface of the skull which was then aligned using bregma and lambda as vertical references. For virus surgeries, viruses were injected at a rate of 100nl/min with a 35-gauge microinjection syringe (World Precision Instruments) connected to a microsyringe pump (UMP3 UltraMicroPump, World Precision Instruments). After viral infusion, the needle was kept at the injection site for a minimum of 5mins and then slowly withdrawn. After surgery, animals were allowed to recover on a heated pad until ambulatory.

For LFP only surgeries, 75μm tungsten wire electrodes (California Fine Wire) were inserted into the BLA (−1.5 AP, −3.3ML, −4.7 DV) and the vHC (−3.2 AP, −3.2 ML, −4.1 DV). A reference and ground screw was implanted over the cerebellum. Electrodes were initially fixed in place with light activated cement (Ivoclar Vivadent) and then affixed to the skull with Metabond (Parkell). Electrodes and the ground wire were then connected to an electrode interface board (EIB-16, Neuralynx) which was then fixed in place using dental cement.

For TEMPO surgeries, a 2:1 mixture of AAV1-CAG-DIO-Ace2N-4AA-mNeon (Virovek) and AAV2-Synapsin-tdTomato (1.23 × 1012 vg ml−1; SignaGen Laboratories) were injected at the following coordinates: for BLA, −1.5 AP, −3.3 ML, −4.8 and −4.6 DV (150μl per site). For vHC, −3.8 AP, − 3.2 ML, −4.1 and −4 DV (250μl per site) at an 11-degree angle. After virus injection, a custom made optrode was implanted in both the BLA and vHC. Each optrode consisted of a 400/430-μm (core/outer) diameter, NA = 0.48, fiber implant (Doric Lenses) with a tungsten LFP wire affixed such that the tip of the electrode protruded 200-300μm beyond the end of the optic fiber. Behavior was performed at least 5 weeks after injection to allow sufficient time for virus expression.

For channelrhodopsin surgeries, mice were bilaterally injected in both the BLA (−1.5 AP, −3.3 ML, −4.8 and −4.6 DV; 150μl per site) and the vHC (−3.8 AP, − 3.2 ML, −4.1 and −4 DV; 250μl per site at an 11-degree angle) with AAV5-EF1α-DIO-ChR2-eYFP (Addgene) to selectively target neuronal populations expressing Cre. After injection of virus, a 200/240-μm (core/outer) diameter, NA = 0.22, was implanted bilaterally in both the BLA and vHC. Behavior was performed at least 4 weeks after injection to allow sufficient time for virus expression.

### Behavioral assays

#### General

Mice were group housed in reversed 12-hr light/dark cycles and all experiments were performed during the dark portion of the cycle. After sufficient time for surgical recovery and/or viral expression, mice underwent multiple rounds of habituation. Each day, mice were brought to the behavioral testing room and transferred to individual cages. These cages were used throughout habituation and testing and acted as individual ‘home’ cages. Mice were first habituated to the behavioral testing room for 30 mins prior to beginning of any further handling each day. Mice were habituated to touch with 2-3 days of handling for 5 mins each day followed by 2 days of habituation to any required tethers for 10-15 mins.

#### Elevated zero maze

For LFP only experiments, mice were first were first recorded in their homecage for 30 mins. Immediately following this, mice were transferred to the elevated zero maze (EZM) for 5 mins. Mice were placed in the entrance of a closed arm, facing into the closed arm. Animals were tested 3 times with a 48hr inter-trial interval.

#### Elevated plus maze

For LFP only and TEMPO experiments, mice were assessed using the elevated plus maze (EPM) for 15 mins. Animals were placed in the center of the EPM facing a closed arm. For channelrhodopsin experiments, EPM sessions lasted 9 mins. In-phase or out-of-phase channelrhodopsin stimulation (473nm, 3-5mW peak power) were delivered during the second and third three minute epochs and were counterbalanced across animals. For in-phase stimulation, a function generator (Agilent 33500B Series Waveform Generator) connected to two lasers generates simultaneous 20Hz trains of 5ms pulses which are coupled to fiber optic implants through a 200μm diameter dual fiber optic patchcord with a guiding socket (Doric Lenses). For out-of-phase stimulation, each 473nm laser was connected to a separate function generator. The triggering of these function generators was offset by 25ms, producing a phase difference of 180°. Real-time light delivery was based on location within the apparatus, with the stimulation zone restricted to the center zone and the proximal third of the closed and open arms.

#### Cross maze

Experiments were performed with the same experimental design as described for EPM with optogenetic stimulation but with a cross shaped maze consisting of 4 closed arms. Animals were placed in the center of the cross maze facing a closed arm on the axis that was classified as ‘Closed 1’. ‘Closed 2’ arms were classified as the two arms that run perpendicular that axis, which are the geographical equivalent of the open arms in the EPM.

### LFP recordings

#### Recordings and analysis

LFP data were sampled at 2KHz and band-pass filtered from 0.1-200Hz using an Open Ephys acquisition board (Open Ephys) connected to an Intan RHD 16-channel headstage (Intan). Electrode placements were verified histologically. Analysis of LFP data was performed using custom MATLAB code. Data was assessed in the following frequency bands: theta band (4-12Hz), beta band (13-30Hz), low gamma band (30-50Hz) and high-gamma band (65-90Hz). Coherence was calculated in 10s non-overlapping windows using MATLAB’s *mscohere* function. Variance of coherence was calculated with a sliding 60s window. For β-burst analysis, multi-taper time-frequency coherence measures were generated with *cohgramc* function from the Chronux toolbox (http://chronux.org). β-bursts were classified as time and frequency-continuous periods of coherence greater than 80% maximum coherence. For EPM runs, β-burst probability was assessed in 500ms windows from 5s before to 5s after the animal exited the center zone. Data was z-scored to probability across the full recording session.

### TEMPO recordings

#### Optical apparatus

A fiber-optic stub (400-μm core, NA = 0.48, low-autofluorescence fiber; Doric Lenses, MFC_400/430-0.48_2.8mm_ZF1.25_FLT) was stereotaxically implanted in each targeted brain region. A matching fiber-optic patch cord (Doric Lenses, MFP_400/430/1100-0.48_2m_FC-ZF1.25) provided a light path between the animal and a miniature, permanently aligned optical bench, or ‘mini-cube’ (Doric Lenses, FMC5_E1(460-490)_F1(500-540)_E2(555-570)_F2(580-680)_S). A single fiber was used to both deliver excitation light to and collect emitted fluorescence from each recording site. The far end of the patch cord and each 1.25-mm-diameter zirconia optical implant ferrule were cleaned with isopropanol before each recording and then securely attached via a zirconia sleeve.

The mini-cube optics allow for the simultaneous monitoring of two spectrally separated fluorophores, with dichroic mirrors and cleanup filters chosen to match the excitation and emission spectra of the voltage sensor and reference fluorophores in use (‘mNeon’ voltage sensor channel: excitation, 460– 490 nm, emission, 500-540 nm; ‘Red’ control fluorophore: excitation, 555–570 nm, emission, 580– 680 nm). The mini-cube optics were sealed and permanently aligned, and all five ports (sample to animal, two excitation lines and two emission lines) were provided with matched coupling optics and FC connectors to allow for a modular system design.

Excitation light for each of the two color channels was provided by a fiber-coupled LED (center wavelengths, 490 nm and 565 nm, Thorlabs, M490F3 and M565F3) connected to the mini-cube by a patch cord (200 μm, NA = 0.39; Thorlabs, M75L01). Using a smaller diameter for this patch cord than for the patch cord from the cube to the animal is critical to reduce the excitation spot size on the output fiber face and thus avoid cladding autofluorescence. LEDs were controlled by a four-channel, 10-kHz-bandwidth current source (Thorlabs, DC4104). LED current was adjusted to give a final light power at the animal (averaged during modulation, see below) of approximately 200 μW for the mNeon channel (460– 490-nm excitation) and 100 μW for the Red channel (555–570-nm excitation).

Each of the two emission ports on the mini-cube was connected to an adjustable-gain photoreceiver (Femto, OE-200-Si-FC; bandwidth set to 7 kHz, AC coupled, ‘Low’ gain of ∼5 × 107 V/W) using a large-core high-NA fiber to maximize throughput (600-μm core, NA = 0.48 (Doric lenses, MFP_600/630/LWMJ-0.48_0.5m_FC-FC).

Note that, for dual-site recordings, two completely independent optical setups were employed, with separate implants, patch cords, mini-cubes, LEDs, photoreceivers and lock-in amplifiers.

### Modulation and lock-in detection

At each recording site, each of the two LEDs was sinusoidally modulated at a distinct carrier frequency to reduce crosstalk due to overlap in fluorophore spectra. The corresponding photoreceiver outputs were then demodulated using lock-in amplification techniques. A single instrument (Stanford Research Systems, SR860) was used to generate the modulation waveform for each LED and to demodulate the photoreceiver output at the carrier frequency. To further reduce crosstalk between recording sites, distinct carrier frequencies were used across at each site (mNeon site 1: 2 kHz; mNeon site 2: 2.5 kHz; Red site 1: 3.5 kHz; Red site 2: 4 kHz). Low-pass filters on the lock-in amplifiers were selected to reject noise above the frequencies under study (cascade of four Gaussian FIR filters with 84-Hz equivalent noise bandwidth; final attenuation of signals were approximately −1 dB (89% of original magnitude) at 20 Hz, −3 dB (71% of original magnitude) at 40 Hz and −6 dB (50% of original magnitude) at 60 Hz).

### Recording

Analog signals were digitized by a multichannel real-time signal processor (Tucker-Davis Technologies, RX-8). The commercial software Synapse (Tucker-Davis Technologies) running on a PC was used to control the signal processor and write data streams to disk. Lock-in amplifier outputs were digitized at 3 kHz. Signals were downsampled to 2KHz for further analysis.

### Analysis

Analysis of TEMPO signals was performed using the signal processing toolbox and custom written MATLAB code. All four signals from the from recording sessions (vHC mNeon, vHC tdTomato, BLA mNeon, BLA tdTomato) were filtered around the frequency of interest using the Matlab function *filitfilt*. mNeon signals for each region were corrected by subtraction of the robust linear regression model (Matlab function *robustfit*) of the tdTomato and mNeon signals across 1s windows. For correlation across full sessions or in zones of the EPM, zero-phase lag correlation was calculated over 250ms non-overlapping windows. For time-locked analysis during EPM zone transitions, 80% overlapping 1s windows were analyzed from 5s before to 5s after the animal exited the center zone. For assessment of TEMPO signal synchrony during β-bursts, LFP signals were first analyzed as described to identify the onset of β-bursts. Time-locked TEMPO signal analysis was then performed from 500ms before β-bursts onset to 500ms after burst onset using a correlation window of 250ms with 80% overlap. Analysis was restricted to β-bursts with a duration of at least 250ms. Data correlations were compared to 100 correlations from randomly shuffled data.

### Histology and imaging

All mice used for behavioral and imaging experiments were anesthetized with Euthasol and transcardially perfused with ice-cold 0.01 M PBS followed by ice-cold 4% paraformaldehyde (PFA) in PBS. Brains were extracted and stored in 4% PFA for 24 h at 4 °C before being stored in PBS. Next, 75μm slices were obtained on a Leica VT100S and mounted on slides using aqueous mounting medium containing DAPI (Vectashield Plust Antifade mounting medium, Vector labs). Sections were imaged using a BZ-X fluorescence microscope (Keyence).

### Human recordings

Intracranial electroencephalography (iEEG) recordings were obtained from 13 human subjects (n=9 Male, n=4 Female) with treatment-resistant epilepsy with surgically implanted semi-chronic intracranial electrodes (for sites included in this study: Ad-Tech 4-contact strip or depth, 10mm center-to-center spacing, 2.3mm exposed diameter; Ad-Tech 10-contact depth, 6 or 5mm center-to-center spacing) for the clinical purpose of seizure localization. All procedures were approved by the University of California, San Francisco Institutional Review Board. All subjects gave written informed consent to participate in the study before surgery. 6 subjects had left hemisphere implantations, 7 had right hemisphere implantations). Subjects’ psychological state was evaluated up to four times daily during their hospitalization period prompted by a research assistant, using a tablet-based, custom-designed questionnaire called the Immediate Mood Scaler (Nahum et al., 2017) (IMS). Details of psychological testing, electrophysiologic recording, data processing, and coherence calculations were reported previously (Kirkby et al 2018).

AMY-HPC beta (β, 13–30 Hz) coherence was calculated in 5 second bins with a 0.5 second sliding window over a 20 minute period centered on an IMS score. Coherence variance was calculated as the mean of the rolling variance (bin = 10) over the mood score window. β-bursts were defined as beginning when coherence increases by a set magnitude (50% of the max coherence during the 20 minute IMS window) above the mean coherence of a 30 second trailing window (e.g. mean coherence during the previous 30 seconds). β-bursts end when coherence drops below 50% of the max coherence. β-bursts were quantified for each time window by summing the area under the curve (AUC) for all bursts, counting the number of bursts, and finding the mean burst duration.

Statistics were performed using the SciPy stats package. Linear regressions between IMS and coherence variance, coherence mean, and sum AUC were performed. As was a linear regression for sum AUC and coherence variance. Student’s t-tests were performed to compare the sum AUC, number of bursts, and burst duration between the top and bottom quartile of IMS scores.

### Statistics

Statistics were calculated using MATLAB code or Graphpad Prism. All statistical parameters, including test statistics, corrections for multiple comparisons and sampling are stated in the main text. Data was tested for normality with the Shapiro-Wilk test. All tests are two-sided and error bars represent standard error of the mean. Sample sizes were based on prior studies. All tests are summarized in Supplemental Table 1.

**Figure S1.**
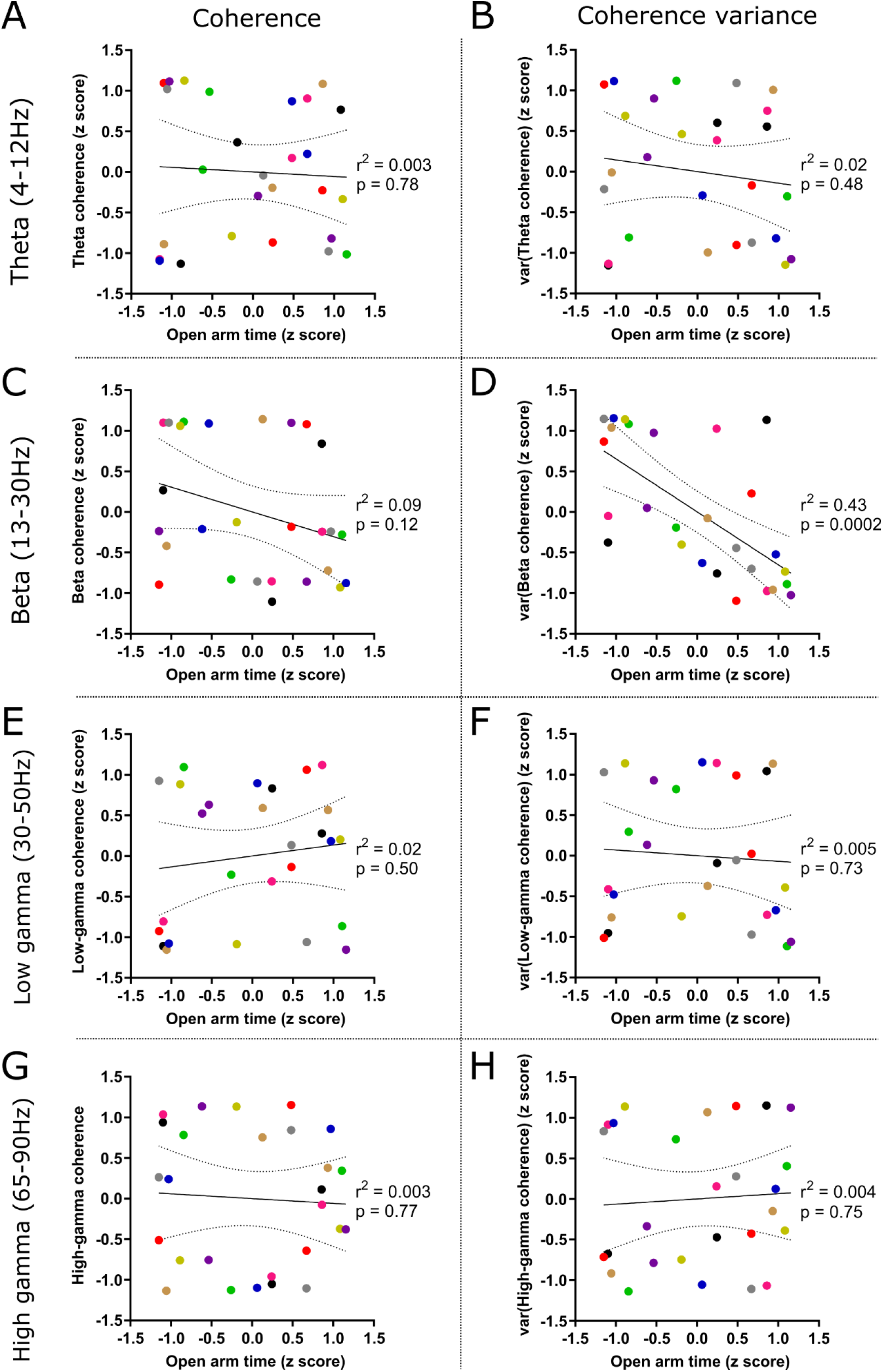
Relationship of EZM open arm time with other homecage LFP metrics. (A) Linear regression of EZM open arm time against homecage LFP mean vHC-BLA theta coherence. Dotted boundaries represent 95% confidence on the regression estimate. (B) Linear regression of EZM open arm time against homecage LFP variance of vHC-BLA theta coherence. Dotted boundaries represent 95% confidence on the regression estimate. (C) Linear regression of EZM open arm time against homecage LFP mean vHC-BLA beta coherence. Dotted boundaries represent 95% confidence on the regression estimate. (D) Linear regression of EZM open arm time against homecage LFP variance of vHC-BLA beta coherence. Dotted boundaries represent 95% confidence on the regression estimate. (E) Linear regression of EZM open arm time against homecage LFP mean vHC-BLA low-gamma coherence. Dotted boundaries represent 95% confidence on the regression estimate. (F) Linear regression of EZM open arm time against homecage LFP variance of vHC-BLA low-gamma coherence. Dotted boundaries represent 95% confidence on the regression estimate. (G) Linear regression of EZM open arm time against homecage LFP mean vHC-BLA high-gamma coherence. Dotted boundaries represent 95% confidence on the regression estimate. (H) Linear regression of EZM open arm time against homecage LFP variance of vHC-BLA high-gamma coherence. Dotted boundaries represent 95% confidence on the regression estimate.

**Figure S2.**
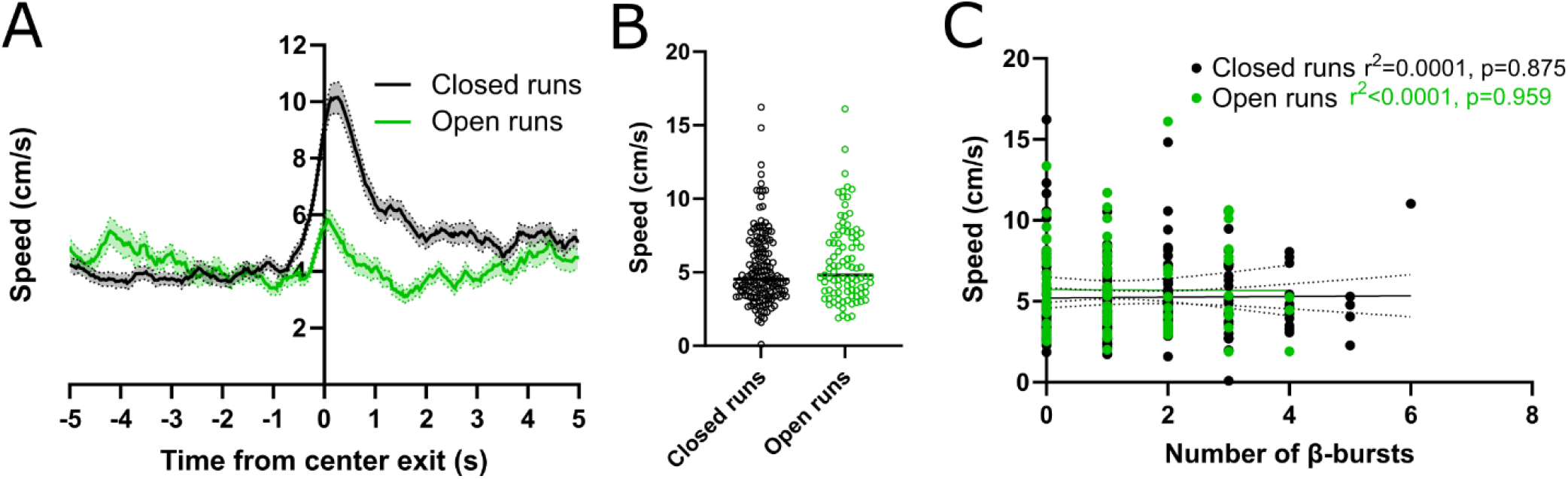
The frequency of occurrence of β-bursts is unaffected by locomotion speed. (A) Locomotion speed during transitions from the EPM center to the closed and open arms (B) Mean speed in the center of the EPM prior to entering the closed or open arm is similar (p=0.2126, closed runs n=166, open runs n=93; Mann-Whitney test). (C) Linear regression between mean locomotion speed and the number of β-bursts for both closed and open arm runs.

**Figure S3.**
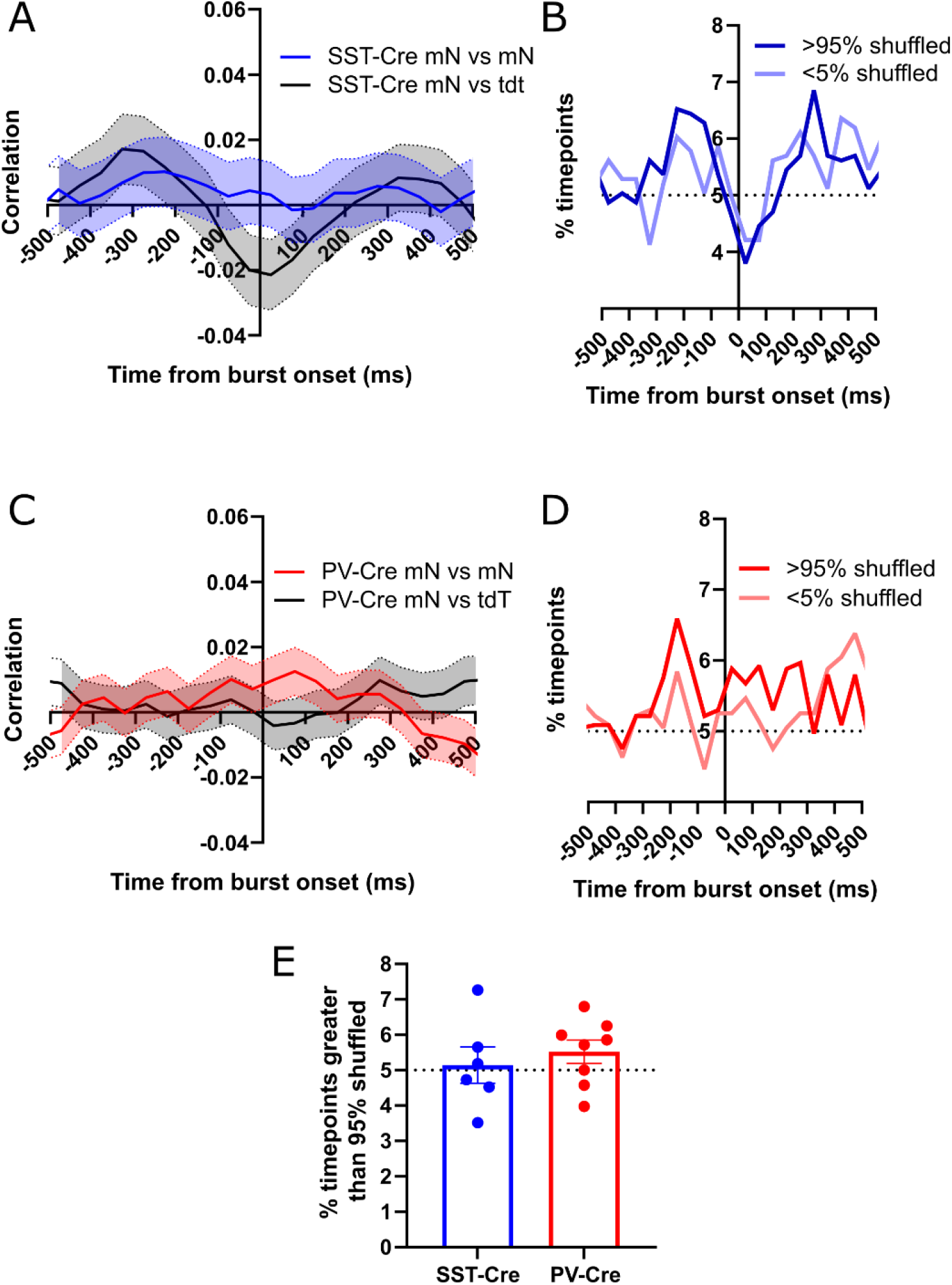
30-50Hz synchrony of SST+ or PV+ interneurons during vHC-BLA β-bursts. (A) SST+ interneuron 30-50Hz synchrony (mNeon correlation values) aligned to the onset of vHC-BLA β-bursts. (B) Time course of SST+ interneuron mNeon correlation values relative to 95% and 5% of shuffled mNeon signals. (C) PV+ interneuron 30-50Hz synchrony (mNeon correlation values) aligned to the onset of vHC-BLA β-bursts. (D) Time course of PV+ interneuron mNeon correlation values relative to 95% and 5% of shuffled mNeon signals. (E) SST+ interneurons and PV+ interneurons exhibit similar 30-50Hz synchrony during vHC-BLA β-bursts (p=0.531, *SST-Cre* n=6, *PV-Cre* n=8; unpaired t test).

**Figure S4.**
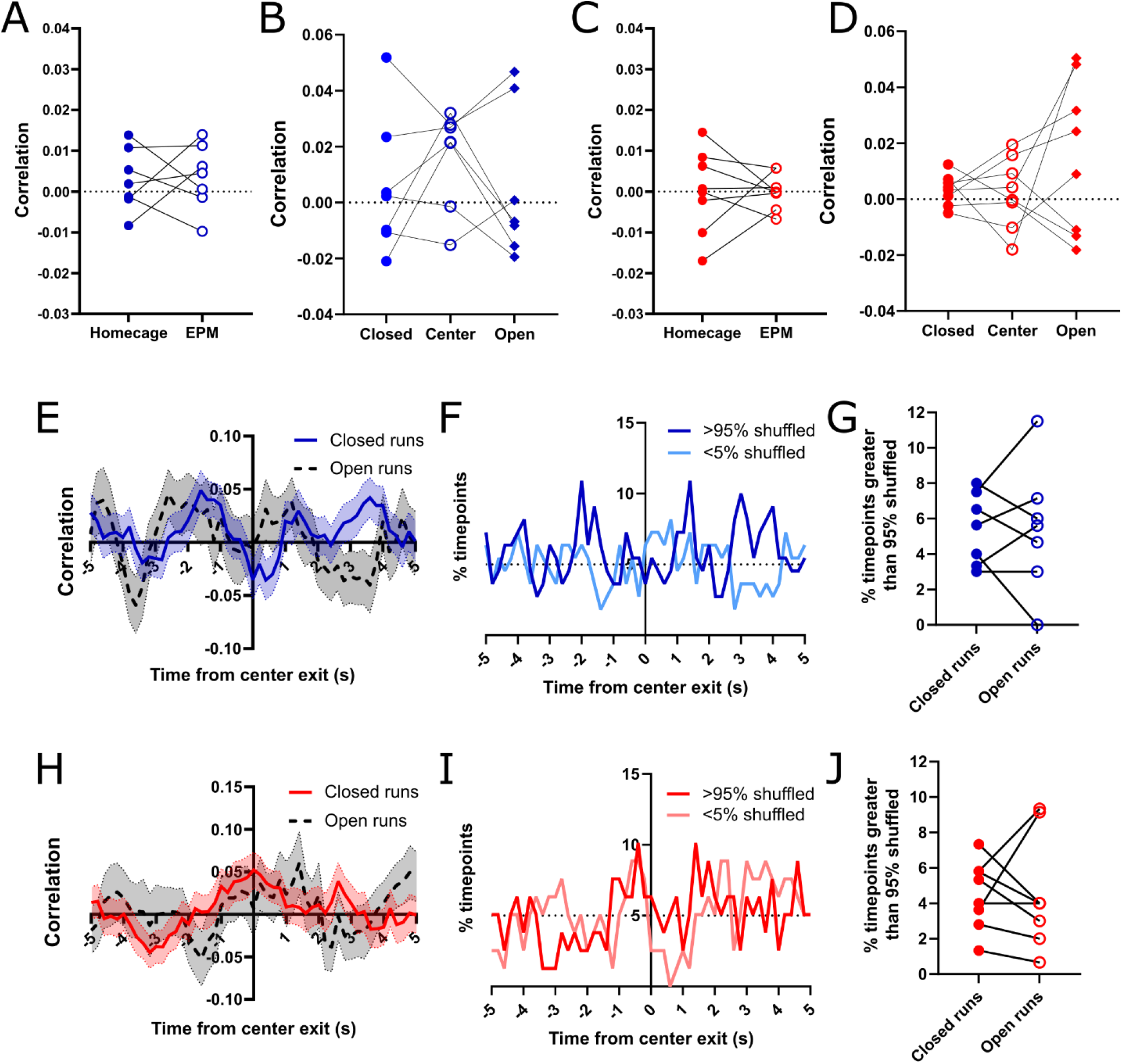
Interneuron 30-50Hz synchrony during EPM behavior. (A) SST+ interneuron 30-50Hz synchrony (mNeon correlation) was similar in EPM sessions and homecage recordings (p=0.8732, n=7 mice; paired t test). (B) SST+ interneuron 30-50Hz synchrony was similar across different zones of the EPM (p = 0.3965, n=7 mice; Repeated measures one-way ANOVA). (C) PV+ interneuron 30-50Hz synchrony (mNeon correlation) was comparable similar in EPM sessions and homecage recordings (p=0.9979, n=8 mice; paired t test). (D) PV+ interneuron 30-50Hz synchrony was similar across different zones of the EPM (p = 0.3062, n=8 mice; Repeated measures one-way ANOVA). (E) SST+ interneuron 30-50Hz synchrony (mNeon correlation values) aligned to closed or open arm entries. (F) Time course the fraction of SST+ interneuron mNeon correlation values (for 30-50Hz synchrony) exceeding / below 95% / 5% of values based on shuffled mNeon signals, during runs into the closed arms. (G) SST+ interneuron synchrony was similar during the 5s prior to entries to the closed or open arms (p=0.991, n=7 mice; paired t test). (H) PV+ interneuron 30-50Hz synchrony (mNeon correlation values) aligned to entries to the closed or open arms. (I) Time course the fraction of PV+ interneuron mNeon correlation values (for 30-50Hz synchrony) exceeding / below 95% / 5% of values based on shuffled mNeon signals, during runs into the closed arms. (J) PV+ interneuron synchrony was similar during the 5s prior to entries to the closed or open arms (p=0.99, n=8 mice; paired t test).

**Figure S5.**
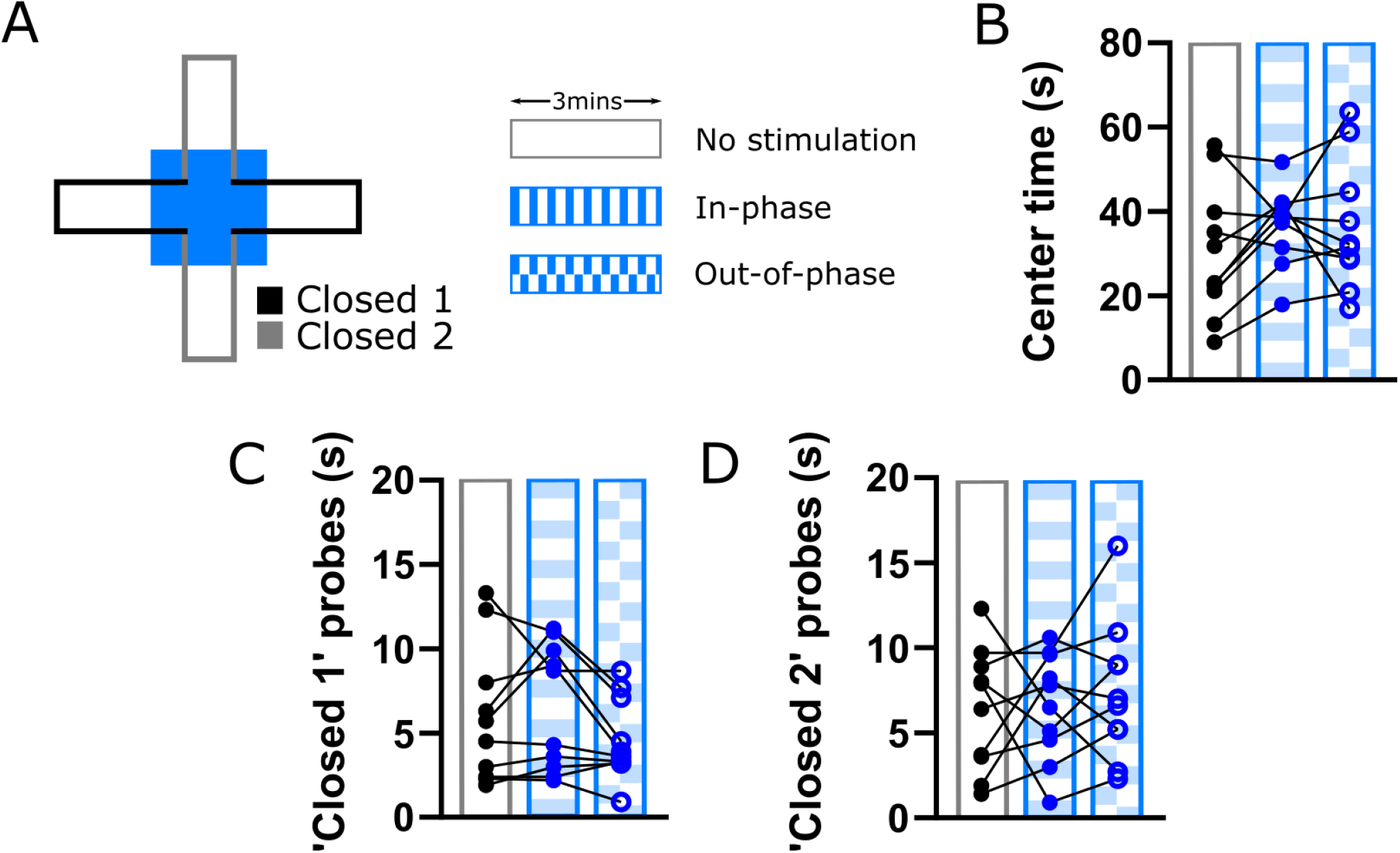
Effects of modulating vHC-BLA β-synchrony on cross-maze behavior. (A) Experimental design: 473nm stimulation was restricted to the center zone and proximal 1/3 of each arm of a cross-maze. Stimulation varied across three 3min epochs: no stimulation, in-phase stimulation and out-of-phase stimulation. (B) In-phase or out-of-phase stimulation at 20Hz had no effect on time in the center of the cross maze (p=0.537; n=10 mice; One-way ANOVA). (C) In-phase or out-of-phase stimulation at 20Hz had no effect on probes into the ‘closed 1’ arms (the geographical equivalent of the EPM closed arms) (p=0.809; n=10 mice; One-way ANOVA). (D) In-phase or out-of-phase stimulation at 20Hz had no effect on probes into the ‘closed 2’ arms (the geographical equivalent of the EPM open arms) (p=0.466; n=10 mice; One-way ANOVA).

---

**Supplemental Table 1.** Details of statistical tests.

| Figure | Data | Test | Result | N | n |
| --- | --- | --- | --- | --- | --- |
| Figure 1B | vHC-BLA β-coherence variance v EZM open arm time | Linear regression | p=0.0002 | N=9 mice | n=27 trials |
| Figure 1D | β-burst frequency v mean vHC-BLA coherence | Linear regression | p<0.0001 | N=9 mice | n=3780 recording windows |
| Figure 1F | High variance/Low variance βcoherence | One sample t-test | p<0.001 | N=9 mice | n=27 trials |
| Figure 1F | High variance/Low variance β-burst frequency | One sample t-test | p=0.0016 | N=9 mice | n=27 trials |
| Figure 1F | High variance/Low variance β-burst duration | One sample t-test | p=0.0028 | N=9 mice | n=27 trials |
| Figure 1F | High variance/Low variance β-burst bandwidth | One sample t-test | p=0.0225 | N=9 mice | n=27 trials |
| Figure 1H | AMY-HPC β-coherence variance v β-burst frequency | Linear regression | p=8.5x10<sup>-14</sup> | N=13 patients | n=110 trials |
| Figure 1I | IMS Score v β-burst frequency | Linear regression | p=4.5x10<sup>-14</sup> | N=13 patients | n=110 trials |
| Figure 1J | Low IMS v High IMS β-burst count | Mann-Whitney test | p<0.0001 | N=13 patients | n=28 trials |
| Figure 2C | Closed v Open run β-burst frequency | Paired t-test | p=0.0122 | N=9 mice | n=9 mice |
| Figure 3I | SST-Cre v PV-Cre β-burst 13-30Hz, 0-250ms post onset, % timepoints >95% shuffled data | Mann-Whitney test | p=0.0127 | SST-cre: N=6 mice, PV-cre: N=8 mice | SST-cre: n=6 mice, PV-cre: n=8 mice |
| Figure 4A | SST-Cre mNeon v mNeon 13-30Hz correlation Homecage v EPM | Paired t-test | p=0.0375 | SST-cre: N=7 mice | SST-cre: n=7 mice |
| Figure 4B | SST-Cre mNeon v mNeon 13-30Hz correlation EPM Closed v Center v Open | Friedman test with Dunn's multiple comparison | Friedman p=0.0207; Closed v Center p=0.0485; Closed v Open p>0.999; Center v Open p=0.0485 | SST-cre: N=7 mice | SST-cre: n=7 mice |
| Figure 4C | PV-Cre 13-30Hz mNeon v mNeon correlation Homecage v EPM | Paired t-test | p=0.2985 | PV-cre: N=8 mice | PV-cre: n=8 mice |
| Figure 4D | PV-Cre 13-30Hz mNeon v mNeon correlation EPM Closed v Center v Open | Repeated measure one-way ANOVA | p=0.7829 | PV-cre: N=8 mice | PV-cre: n=8 mice |
| Figure 4H | SST-Cre 13-30Hz Closed v Open runs, 0-5s prior to center exit, timepoints >95% shuffled data | Paired t-test | p=0.039 | SST-cre: N=7 mice | SST-cre: n=7 mice |
| Figure 4K | PV-Cre 13-30Hz Closed v Open runs, 0-5s prior to center exit, timepoints >95% shuffled data | Paired t-test | p=0.3557 | PV-cre: N=8 mice | PV-cre: n=8 mice |
| Figure 4O | SST-Cre v PV-Cre 13-30Hz open arm probes, 0-1.5s pre probe end, % timepoints >95% shuffled data | Mann-Whitney test | p=0.0267 | SST-cre: N=7 mice, PV-cre: N=8 mice | SST-cre: n=7 mice, PV-cre: n=8 mice |
| Figure 5C | SST-Cre 20Hz, SST-Cre 8Hz and PV-Cre 20Hz no stimulation v in-phase v out-of-phase Center time | Two-way ANOVA with Tukey's multiple comparison | *Frequency x Session:*<br>p<0.0001; *SST-Cre 20Hz:*<br>No stim v in-phase p=0.1872, No stim v out-of-phase p=0.009, in-phase v out-of-phase p=0.0365; *SST-Cre 8Hz:* No stim v in-phase p=0.8210, No stim v out-of-phase p=0.2446, in-phase v out-of-phase p=0.1202; *PV-Cre 20Hz:* No stim v in-phase p=0.157, No stim v out-of-phase p=0.2237, in-phase v out-of-phase p=0.9988 | SST-Cre 20Hz: N=10 mice, SST-Cre 8Hz: N=7 mice, PV-Cre 20Hz: N=8 mice | SST-Cre 20Hz: n=10 mice, SST-Cre 8Hz: n=7 mice, PV-Cre 20Hz: n=8 mice |
| Figure 5D | SST-Cre 20Hz, SST-Cre 8Hz and PV-Cre 20Hz no stimulation v in-phase v out-of-phase Open time | Two-way ANOVA with Tukey's multiple comparison | *Frequency x Session:*<br>p=0.6816 | SST-Cre 20Hz: N=10 mice, SST-Cre 8Hz: N=7 mice, PV-Cre 20Hz: N=8 mice | SST-Cre 20Hz: n=10 mice, SST-Cre 8Hz: n=7 mice, PV-Cre 20Hz: n=8 mice |
| Figure 5E | SST-Cre 20Hz, SST-Cre 8Hz and PV-Cre 20Hz no stimulation v in-phase v out-of-phase Closed time | Two-way ANOVA with Tukey's multiple comparison | *Frequency x Session:*<br>$p=0.0219$; *SST-Cre 20Hz:*<br>No stim v in-phase $p=0.6198$, No stim v out-of-phase $p=0.0491$, in-phase v out-of-phase $p=0.3028$;<br>*SST-Cre 8Hz:* No stim v in-phase $p=0.9512$, No stim v out-of-phase $p=0.2085$, in-phase v out-of-phase $p=0.3392$; *PV-Cre 20Hz:* No stim v in-phase $p=0.3646$, No stim v out-of-phase $p=0.1801$, in-phase v out-of-phase $p=0.9021$ | SST-Cre 20Hz: N=10 mice, SST-Cre 8Hz: N=7 mice, PV-Cre 20Hz: N=8 mice | SST-Cre 20Hz: n=10 mice, SST-Cre 8Hz: n=7 mice, PV-Cre 20Hz: n=8 mice |
| Figure 5F | SST-Cre 20Hz, SST-Cre 8Hz and PV-Cre 20Hz no stimulation v in-phase v out-of-phase Open arm probe time | Two-way ANOVA with Tukey's multiple comparison | *Frequency x Session:*<br>$p=0.0016$; *SST-Cre 20Hz:*<br>No stim v in-phase $p=0.7055$, No stim v out-of-phase $p<0.0001$, in-phase v out-of-phase $p=0.0008$;<br>*SST-Cre 8Hz:* No stim v in-phase $p=0.8935$, No stim v out-of-phase $p=0.9691$, in-phase v out-of-phase $p=0.7697$; *PV-Cre 20Hz:* No stim v in-phase $p=0.7009$, No stim v out-of-phase $p=0.6244$, in-phase v out-of-phase $p=0.9916$ | SST-Cre 20Hz: N=10 mice, SST-Cre 8Hz: N=7 mice, PV-Cre 20Hz: N=8 mice | SST-Cre 20Hz: n=10 mice, SST-Cre 8Hz: n=7 mice, PV-Cre 20Hz: n=8 mice |
| Figure 5G | SST-Cre 20Hz, SST-Cre 8Hz and PV-Cre 20Hz no stimulation v in-phase v out-of-phase Closed runs | Two-way ANOVA with Tukey's multiple comparison | *Frequency x Session:*<br>$p=0.0833$; *SST-Cre 20Hz:*<br>No stim v in-phase $p=0.0094$, No stim v out-of-phase $p=0.9461$, in-phase v out-of-phase $p=0.0221$;<br>*SST-Cre 8Hz:* No stim v in-phase $p=0.6703$, No stim v out-of-phase $p=0.5238$, in-phase v out-of-phase $p>0.999$; *PV-Cre 20Hz:* No stim v in-phase $p=0.8674$, No stim v out-of-phase $p=0.7986$, in-phase v out-of-phase $p=0.9620$ | SST-Cre 20Hz: N=10 mice, SST-Cre 8Hz: N=7 mice, PV-Cre 20Hz: N=8 mice | SST-Cre 20Hz: n=10 mice, SST-Cre 8Hz: n=7 mice, PV-Cre 20Hz: n=8 mice |
| Figure 5H | SST-Cre 20Hz, SST-Cre 8Hz and PV-Cre 20Hz no stimulation v in-phase v out-of-phase Open runs | Two-way ANOVA with Tukey's multiple comparison | *Frequency x Session:*<br>p=0.4907 | SST-Cre 20Hz: N=10 mice, SST-Cre 8Hz: N=7 mice, PV-Cre 20Hz: N=8 mice | SST-Cre 20Hz: n=10 mice, SST-Cre 8Hz: n=7 mice, PV-Cre 20Hz: n=8 mice |
| Figure S1A | vHC-BLA theta-coherence v EZM open arm time | Linear regression | p=0.78 | N=9 mice | n=27 trials |
| Figure S1B | vHC-BLA theta-coherence variance v EZM open arm time | Linear regression | p=0.48 | N=9 mice | n=27 trials |
| Figure S1C | vHC-BLA beta-coherence v EZM open arm time | Linear regression | p=0.12 | N=9 mice | n=27 trials |
| Figure S1D | vHC-BLA beta-coherence variance v EZM open arm time | Linear regression | p=0.0002 | N=9 mice | n=27 trials |
| Figure S1E | vHC-BLA low gamma-coherence v EZM open arm time | Linear regression | p=0.5 | N=9 mice | n=27 trials |
| Figure S1F | vHC-BLA low gamma-coherence variance v EZM open arm time | Linear regression | p=0.73 | N=9 mice | n=27 trials |
| Figure S1G | vHC-BLA high gamma-coherence v EZM open arm time | Linear regression | p=0.77 | N=9 mice | n=27 trials |
| Figure S1H | vHC-BLA high gamma-coherence variance v EZM open arm time | Linear regression | p=0.75 | N=9 mice | n=27 trials |
| Figure S2B | Closed v Open run center zone speed | Mann-Whitney test | p=0.2126 | N=9 mice | Closed runs: n=166 runs, Open runs: n=93 runs |
| Figure S2C | Center zone speed v number of β-bursts | Linear regression | Closed runs: p=0.875; Open runs: p=0.959 | N=9 mice | Closed runs: n=166 runs, Open runs: n=93 runs |
| Figure S3E | SST-Cre v PV-Cre β-burst 30-50Hz, 0-250ms post onset, % timepoints >95% shuffled data | Unpaired t-test | p=0.531 | SST-cre: N=6 mice, PV-cre: N=8 mice | SST-cre: n=6 mice, PV-cre: n=8 mice |
| Figure S4A | SST-Cre mNeon v mNeon 30-50Hz correlation Homecage v EPM | Paired t-test | p=0.8048 | SST-cre: N=7 mice | SST-cre: n=7 mice |
| Figure S4B | SST-Cre mNeon v mNeon 30-50Hz correlation EPM Closed v Center v Open | Repeated measure one-way ANOVA | p=0.3965 | SST-cre: N=7 mice | SST-cre: n=7 mice |
| Figure S4C | PV-Cre 30-50Hz mNeon v mNeon correlation Homecage v EPM | Paired t-test | p=0.9979 | PV-cre: N=8 mice | PV-cre: n=8 mice |
| Figure S4D | PV-Cre 30-50Hz mNeon v mNeon correlation EPM Closed v Center v Open | Repeated measure one-way ANOVA | p=0.3062 | PV-cre: N=8 mice | PV-cre: n=8 mice |
| Figure S4G | SST-Cre 30-50Hz Closed v Open runs, 0-5s prior to center exit, timepoints >95% shuffled data | Paired t-test | p=0.991 | SST-cre: N=7 mice | SST-cre: n=7 mice |
| Figure S4J | PV-Cre 30-50Hz Closed v Open runs, 0-5s prior to center exit, timepoints >95% shuffled data | Paired t-test | p=0.99 | PV-cre: N=8 mice | PV-cre: n=8 mice |
| Figure S5B | SST-Cre 20Hz no stimulation v in-phase v out-of-phase Cross maze center time | Two-way ANOVA with Tukey's multiple comparison | *Frequency x Session:*<br>p=0.537 | SST-Cre: N=10 mice | SST-Cre: n=10 mice |
| Figure S5B | SST-Cre 20Hz no stimulation v in-phase v out-of-phase Closed 1 arm head exploration | Two-way ANOVA with Tukey's multiple comparison | *Frequency x Session:*<br>p=0.809 | SST-Cre: N=10 mice | SST-Cre: n=10 mice |
| Figure S5B | SST-Cre 20Hz no stimulation v in-phase v out-of-phase Closed 2 arm head exploration | Two-way ANOVA with Tukey's multiple comparison | *Frequency x Session:*<br>p=0.466 | SST-Cre: N=10 mice | SST-Cre: n=10 mice |

